# β-Amyloid peptides tailor switching behaviors of Donor-Acceptor Stenhouse Adducts

**DOI:** 10.1101/2020.10.04.325696

**Authors:** Chao Zheng, Yue Yu, Kuang Shi, Biyue Zhu, Heng Zhou, Shao-Qing Zhang, Jing Yang, Liang Shi, Chongzhao Ran

## Abstract

Molecular switching plays a critical role in biological and displaying systems. Here we demonstrate the first use of peptides to operate molecular switches of donor-acceptor Stenhouse adducts (DASAs), a series of negative photochromes that are highly promising for applications ranging from smart material to biological systems. Fluorescence imaging proved Aβ40 species could make SHA-2 more stable in the linear configuration than without peptide and decrease the rate of molecular switching. According to molecular dynamics simulation, SHA-2 bound to protein resulted in substantial changes in the tertiary structure of Aβ40 monomer with the region of Glu22-Ala30 partially unfolded and being more exposed to water. This structural change is likely to impede the aggregation of Aβ40, as evidenced by fluorescence and ProteoStat^®^ aggresome detection experiments. SHA-2 is able to inhibit the aggregation of Aβ40 by producing the off-pathway structures. These results open ample opportunities for optically addressable potential widely apply DASAs in the biological system based on this peptides-tailor process.

## Introduction

Molecular switch has great potential to control molecule’s functions, and tailoring the switching behavior can be used to fine tuning the functions in different systems, including biological systems and displaying systems^1^. To tailor the switch behavior, chemical-stimulations, which including electrochemical-stimulation^2^, thermo-stimulation^3^, pH-stimulation^4^, and photo-stimulation^5^, are conventionally used approaches. Photoswitching of E/Z isomers of protonated Schiff base (PSB)-retinal chromophore, green fluorescent protein/phytochrome, guanine nucleotides controlled regulation of G proteins^6^, and allosteric regulation of enzyme active/protein conformation are the important molecular switches in biological processes.^7^ Especially, photoswitching function of peptides is a very relevance research field. For examples, photoregulation of peptide conformation^1, 8^ and photocontrol biological activity of somatostatin by azobenzene unit^9^, and PSB-retinal based molecular switch to photocontrol the secondary structure of a model peptide^10^. However, two intrinsic disadvantages are associated with these approaches, i.e. no significantly larger change in the end-to-end distance (or molecular contraction) upon isomerization^11^ and the need of troublesome high-energy UV light. To address the deleterious UV light problem, photochromic compounds that can be controlled by visible light have been developed^5b, 12^. However, the results are significantly deficit and scientists have largely ignored the reversible photochromism^13^.

The switching behavior of donor-acceptor Stenhouse adducts (DASAs), which is a class of reversible photochromic molecules that had been overlooked for a very long time since it was discovered in 1870 year ^14^. Due to its versatile switch functions by low-energy visible light, research on DASA has been revived since 2014 ^15^. A major advantage over most other photochromic molecules is its facile synthetic access, modularity, and tenability. DASAs that are responsive to visible or NIR light are more attractive than traditional UV-light-responsive molecules such as azobenzene^16^, spiropyran^17^, coumarin^18^, and o-nitrobenzyl molecules^19^. Irradiation with visible or near-infrared light can induce linear-to-cyclic isomerization of DASAs in aromatic solvents, while cyclic-to-linear isomerization arises upon heating in the dark. On the other hand, in protic solvents such as methanol and water, cyclic zwitterionic (ring-closed) form is stabilized under heating/or in the dark, which due to water molecules coordinate with DASAs and stabilize the intermediates and cyclization thermodynamically favors the cyclic isomers.^20^ This process of linear to cyclic switching is irreversible in water except in a water-soluble Pd^II^_8_ molecular vessel.^21^ Remarkably, the molecular contraction from strongly colored triene form to a colorless zwitterionic (ring-closed) form is accompanying with huge difference in wavelength and solubility. A few mechanisms proposed^22^ that, experimentally^23^, computationally^23a, 24^, the hydroxyl group was critical^25^, and water likely involved in the crucial steps of DASA isomerization^20, 25^; however, it has relatively modest influence on overall switching rates in DMSO.^26^ Although the mechanism of this photoswitching process is not totally clear, numerous applications have already been reported in many material fields, such as secret information coding^27^ photo printing^28^, surface patterning^29^, photoresponsive liquid crystals^30^, sensors^31^, second-order nonlinear optical properties^32^, and the functionalization of magnetic nanoparticles^33^. It has also been applied to drug release^34^, and compatibility with orthogonal switches^28, 35^ and other biological and medicinal applications such as photopharmacology attracting tremendous interest^1, 36^. Despite its wide applications in material fields, to the best of our knowledge, there is no research on whether peptides can tailor the isomerization of DASAs, and such studies are critical for the understanding of DASA switch mechanism and its applications in biomedicine.

Light irradiation is the most used approach to manipulate the switching; however, light stimulation is not mandatory for all cases. For example, in polar protic solvents (e.g. water and methanol),^37^ a background reaction from linear (**L**) to cyclic (**C**) in the dark is observed by Feringa’s group^22–23, 25^, where the compound undergoes cyclization independently of light. This opens a new window for biology transformation in physiological environment. In our previous studies, we have designed several near-infrared fluorescent imaging probes based on hydrophilic and hydrophobic properties of β-amyloid (Aβ) species^38^. We hypothesized that hydrophobic amino acids of Aβ peptide could bind to the colored linear triene form (**A1** to **A3**) of DASAs, and this hydrophobic environment could slow down DASA switching from the colored linear triene form (**A**) to the colorless cyclic form (**B**). In this report, we demonstrated β-Amyloid (Aβ) peptides could be used to tailor the switching behavior of DASAs. Meanwhile, this binding process could be used to suppress the aggregation of Aβ peptides.

## Results

### 1. β-Amyloid monomer tailors switching behaviors of SHA-2 (Fluorescence and Kinetics modeling)

Infrared and ultraviolet absorption spectra of three generations of DASA compounds have been investigated so far, however, no fluorescence spectra of these types of compounds have been reported. To obtain the DASA compound falling into the near infrared wavelength range, we employed the standard DASA synthetic route, in which hydroxypyridone was used as the acceptor to fuse with furfural by the Knoevenagel condensation, and then 2-methylindoline was selected as the donor moiety^39^. With this method, the third generation DASAs compound SHA-2 could be easily obtained (Figure 2a). Alaniz et al reported that many absorption spectral peaks (*λ*_ex_) for the third generation DASAs with 2-methylindoline donor were longer than 600 nm^39^. Therefore, it was reasonable to speculate that, the fluorescence emission wavelength of SHA-2 will fall in the near-infrared window. The excitation of SHA-2 in dichloromethane is 675nm, which is assigned to **A**. However, the 675nm peak gradually disappeared and a new peak at 325nm, which could be assigned to **B,** was observed when SHA-2 was incubated in PBS buffer. When it was excited with 325nm, we could observe a 379nm peak that could be assigned to configuration **B** (Figure 2b). Meanwhile, the emission peak of SHA-2 in dichloromethane is 719nm, which could be ascribed to configuration **A.** We found that the emission peak of SHA-2 in a water solution was 680 nm, which could be ascribed to **A1**. Notably, the fluorescence intensity diminished rapidly within ten minutes. Moreover, the emission peak was red shifted gradually from 680 nm to 720 nm (Figure 2c). These phenomena are consistent with Feringa’s recent report^22–23^ and our DFT calculations. Our DFT calculations indicate that the red shift from 680 nm to 720 nm is likely due to the isomerization of **A1** to **A2** or **A3**, which have similar emission wavelengths (see Supporting Information). In addition, the fluorescence intensity continued to decrease until it was stabilized in this process. The phenomena of fluorescence experiment also indicated that **A1** could quickly convert to the close form **B** through **A2** and **A3**, which were defined as intermediate, in aqueous solution. We first verified the isomerization process with fluorescence.

**Figure 1.**
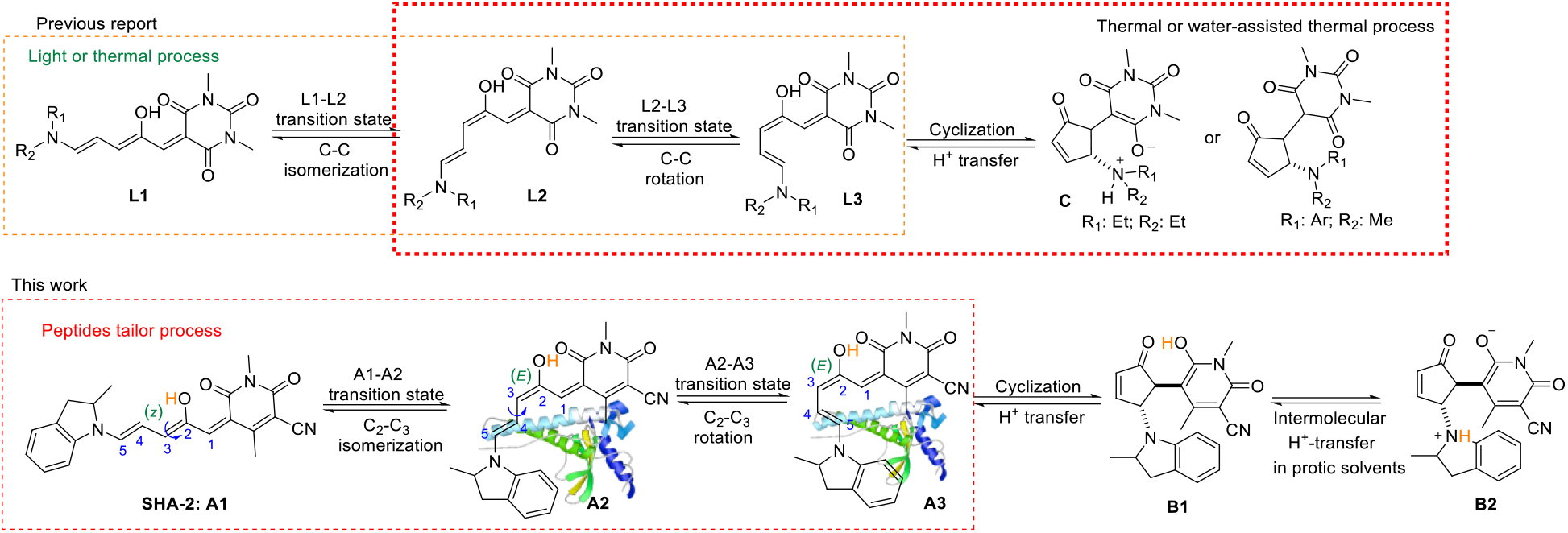
Hypothesis of Aβ peptide tailoring of molecular switching of DASAs (SHA-2)

**Figure 2.**
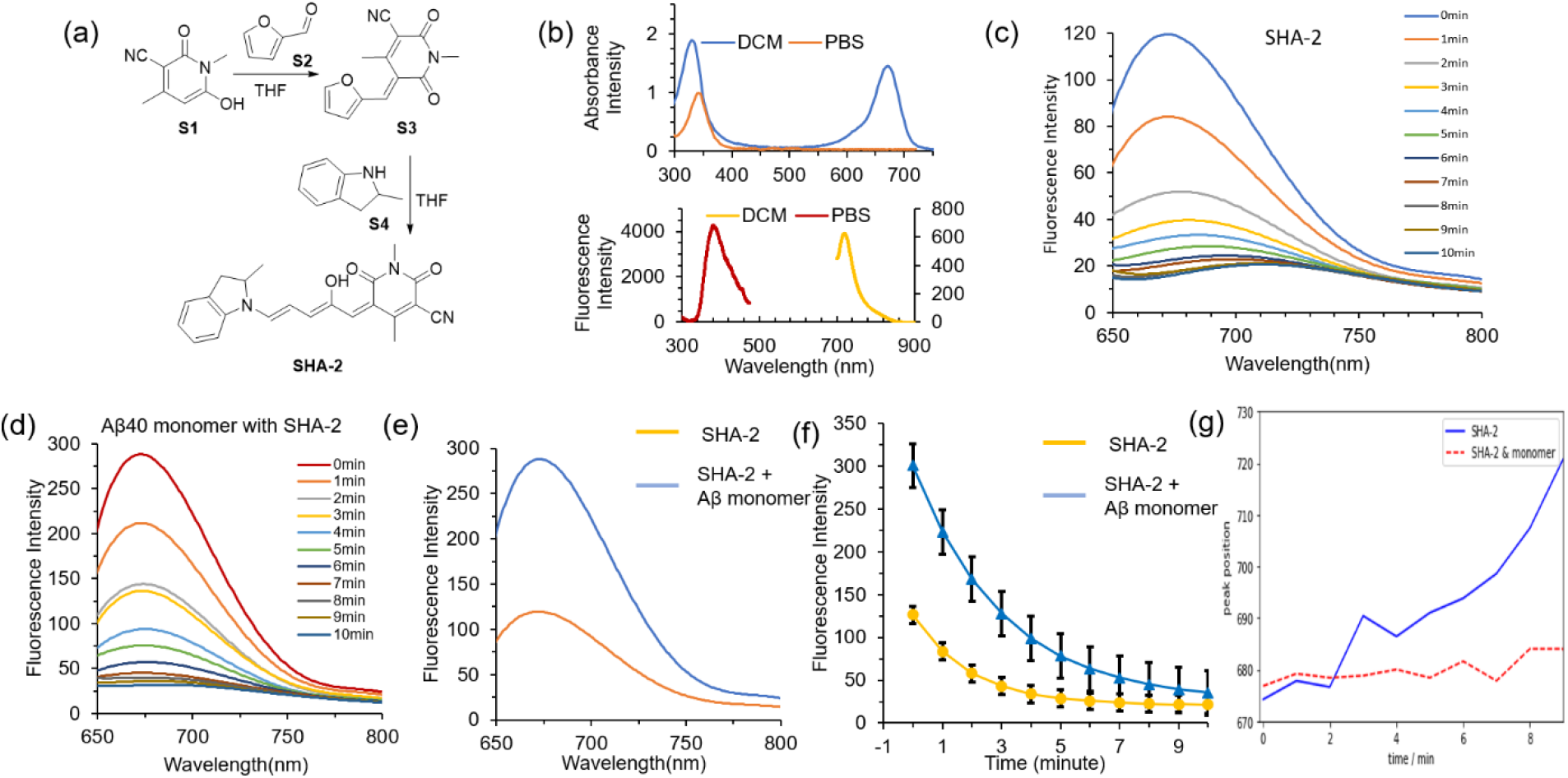
(**a**) The synthetic route of SHA-2; (**b**) Absorption and fluorescent of SHA-2; (**c**) Fluorescence of SHA-2 between 10 min in PBS buffer; (**d**) Fluorescence of SHA-2 between 10 min in Aβ40 monomer PBS solution; (**e**) Fluorescence of SHA-2 at 0 min with or without Aβ40 monomer solution; (**f**) Kinetics fitting results for different SHA-2 solution; (**g**) red-shift ratio for different SHA-2 solution.

According to previous reports, cyclic DASAs **B** are more stable when water exists^15b^, and the linear-to-cyclic switching even occurs under dark in aqueous environment. Considering this is a thermodynamically stable process, it is considerably difficult to stabilize SHA-2 in a particular linear conformation (**A1**, **A2** or **A3**)^22–23, 25^. In this report, we hypothesized that Aβ40 peptide could tailor the switching behavior of DASAs through stabilizing the transient state of **A**. We proved this concept with binding affinity and fluorescence intensities of SHA-2 with/without synthetic Aβ40 monomer. When we incubated SHA-2 with Aβ40 monomers, there was an apparent increase of the fluorescent intensity (2-fold) at 680 nm (Figure 2e), suggesting that Aβ40 can bind to the linear forms of SHA-2. However, with the increasing of incubation time, the increased intensity was gradually decreased. From Fig.2c–2d, after 10 minutes, the intensity was similar to the solution without Aβ40 monomers. We assume that the intensity decline at 680 nm means that some of conformation **A** disappears, and speculated that the decreasing rate of fluorescent intensity could be slower in the presence of Aβ40 monomers. Using the decreasing of fluorescent intensity peak of SHA-2 with/without peptides system (Fig.2c–2d) to quantify the rate of intensity change, we performed kinetic modeling based on the dissociation of one-phase exponential decay (the equation and the fitting results are given in Supporting Information). Our data showed that the decay half-life of SHA-2 was 1.350 min in PBS, and 2.083 min with Aβ40 monomer, respectively (Fig. 2f). To investigate whether the presence of Aβ can slow down the red-shift, we monitored the emission peaks at different time-points, and found that the red-shift ratio of SHA-2 is slower when incubated with Aβ40 monomer (Fig.2g).

Taken together, the above results indicate that Aβ40 monomer could make SHA-2 more stable in the configuration **A** than without the peptide and decrease the rate of molecular switching.

### 2. Dynamic binding sites of β-Amyloid peptides tailor SHA-2 switching (Epitope alteration experiment and simulation)

Recently, our group demonstrated that small molecules could alter epitope binding to its antibody, resulting enhanced or weakened binding of the antibody to its antigen.^40^ To investigate whether SHA-2 has this alteration effect, we incubated Aβ monomers with and without SHA-2, and Aβ antibody 4G8 (epitope Leu17-Val24) was used in our dot-blotting assay. From Fig.3a, it is clear that SHA-2 can significantly weaken the binding between Aβ and 4G8 antibody, suggesting that SHA-2 can bind to Aβ peptides. We speculate that SHA-2 can bind well with Aβ peptides, and in return Aβ monomer can tailor this switching behavior.^38^ To verify this, we performed classical molecular dynamics (MD) simulations for three aqueous systems using GROMOS 54a7 force field^41^: SHA-2 in water, Aβ40 monomer in water, and both solvated in water (denoted as Aβ40-SHA-2). For each system, five independent MD simulations were conducted, and each was run for 400 ns after proper equilibrations, making a total of 2 μs simulation time for each system. For the Aβ40-SHA-2 simulations, Aβ40 monomer and SHA-2 were initially solvated independently, allowing binding to occur naturally to avoid bias. The details of MD simulations are provided in Section 6 of the SI. According to MD simulations, four out of five simulations suggested that SHA-2 bound to Aβ40 at similar positions and formed hydrogen bonds with Glu22 (or Asp23) and Lys28. One representative configuration of SHA-2 binding to Aβ40 is shown in Fig. 3(b). The hydroxyl group near tetrahydropyrimidine-trione of SHA-2 served as a hydrogen bond donor to Glu22-Oε. Note that one of the four simulations where SHA-2 bound at the similar position showed that SHA-2 can also possibly serve as the hydrogen bond donor to Asp23-Oδ. The carbonyl oxygen on tetrahydropyrimidine-trione ring served as a hydrogen bond acceptor and formed a hydrogen bond with positive charged Lys28-Nζ. The hydrophilic part of SHA-2 around dioxo-tetrahydropyridine interacted with Glu22 (or Asp23) and Lys28 via hydrogen bonds so that the specific region of SHA-2 was kept stable, and the more hydrophobic region of SHA-2 around indoline locates at an environment stabilized by hydrophobic-aromatic interaction with Phe19 (**Figure 3**). Take together, both the epitope alteration experiment and MD simulation suggest that SHA-2 indeed binds stably with Aβ40.

**Figure 3.**
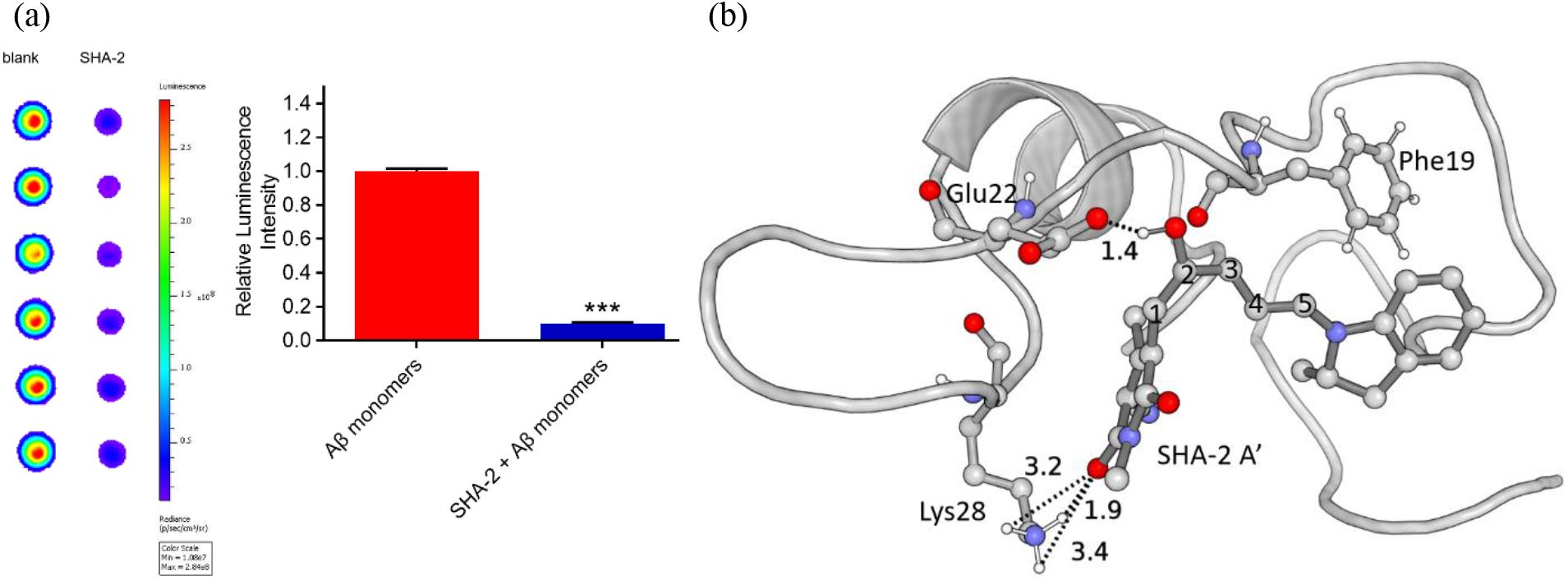
(**a**) Epitope alteration experiment; (**b**) SHA-2 binding position on Aβ40, Glu22 and Lys28 were labelled and possible hydrogen bonds formed between Aβ40 and hydroxypyridone of SHA-2 were shown with dashed lines, and the distances were labelled in Å. Bridge carbon atoms connecting dioxo-tetrahydropyridine and indoline of SHA-2 were numbered from 1 to 5, and the figure showed A2 SHA-2 configuration. Phel9 was also labelled as it potentially formed hydrophobic-aromatic interactions with SHA-2.

In order to elucidate how the binding controls SHA-2 configuration changes, we computed the distributions of the dihedral angles of 1-2-3-4 and 2-3-4-5 in SHA-2 (labels showed in Figure 1) for SHA-2 bound to Aβ40 and SHA-2 solvated in aqueous solution, shown in panels (a) and (b) of Fig. 4, respectively. In all four simulations, Aβ40-SHA-2 systems show similar SHA-2 binding positions on Aβ40 (**Figure 3b**), and during the simulations SHA-2 remained at the binding site. From Fig. 4 the dihedral angles on SHA-2 upon binding to Aβ40 monomer primarily represent **A2** configuration, where dihedral 1-2-3-4 adopts mainly the *cis* configuration and dihedral 2-3-4-5 maintains prominently in the *trans* configuration. In comparison, when SHA-2 is solvated in aqueous solution, all isomers, **A1, A2**, and **A3** were thermally accessible with **A3** being dominant in our simulation. Note that in one of the five independent Aβ40-SHA-2 simulations, SHA-2 weakly binds to the N-terminus of Aβ40, and exposes itself to water, thereby leading to dihedral angle distributions similar to SHA-2 solution without Aβ40. Although different force fields may give different dihedral angle distributions, we expect that **A2** remains the dominant configuration of linear SHA-2 upon binding to Aβ40 monomer as it allows the energetically favorable hydrogen bonding (two hydrogen bonds) and hydrophobic pi-pi interaction between SHA-2 and Phe19 of Aβ40. The binding of SHA-2 to Aβ40 in the form of **A2** stabilizes the open form of SHA-2, impeding its conversion to the closed form.

**Figure 4.**
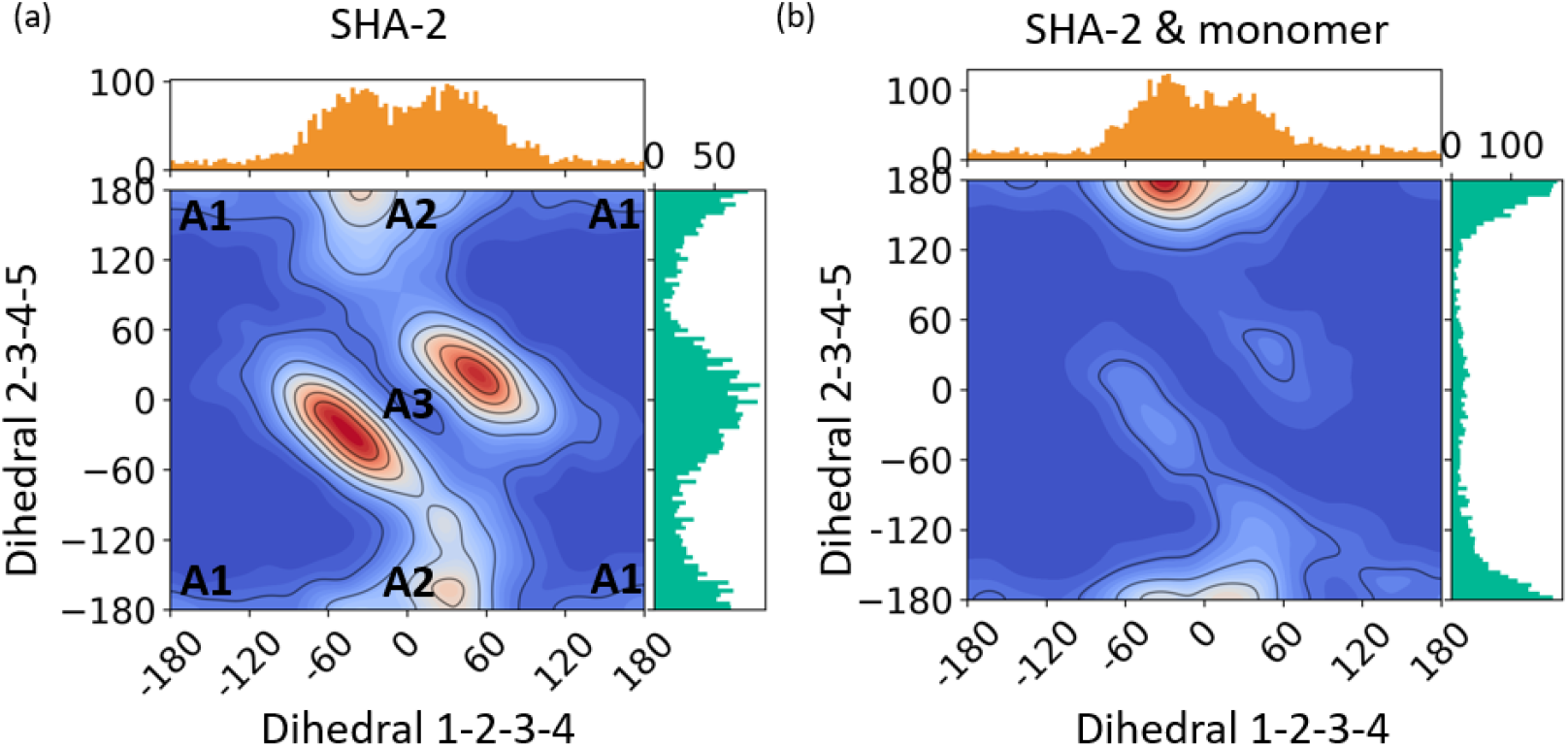
1D and 2D distributions of SHA-2 dihedral angles 1-2-3-4 and 2-3-4-5. (**a**). SHA-2 solvated in water. The typical regions that correspond to different isomers of linear SHA-2 (**A1**, **A2**, and **A3**) are indicated in the 2D distribution. (**b**). SHA-2 bound to Aβ40 monomer.

### 3. Apply Aβ40 species to tailor DASAs switch in tissue level (Plate imaging; Staining slides and light irradiation)

To explore the possibility of more complex protein systems tailor DASAs molecular switching behavior. We expand the peptide system to Aβ40 oligomer and Aβ40 aggregate. When we incubated SHA-2 with Aβ40 aggregate and Aβ40 oligomer, as we expected, there was an apparent increase of the fluorescent intensity, 7-folds and 4-folds, respectively (Figure 5a). Same as Aβ40 monomer, with the increasing of incubation time, the increased intensity was gradually decreased (Figure 5a–5c). As previous for Aβ40 monomer, kinetic modeling studies were performed base on one phase exponential decay, and we found that the decay half-life of SHA-2 was 1.797 minutes with Aβ40 oligomer solutions, and 1.517 minutes with Aβ40 aggregate solutions, respectively. For Aβ40 oligomers, the redshift has been slowed down, while no red shift was observed for Aβ40 aggregates. To confirm that Aβ40 species can change the trajectory of the isomerization of SHA-2 in solutions, we also imaged the solutions on an imaging system (IVIS imaging system) with a 96-well plate. The images were captured at different time point. From Fig.5d, Aβ40 species can significantly delay the disappearance of SHA-2 fluorescence, further suggesting that Aβ40 can slow down the conversion from the A isoforms to the B isoform. To further investigate the interaction between SHA-2 and Aβ species in a biologically relevant environment, we first stained a brain slice from an APP/PS1 transgenic mouse. As show in microscopic images in Figure 5e, SHA-2 could specifically highlight Aβ plaques. To test whether Aβ species can slow down the closing process of SHA-2. We used red and green light to illuminate the slices in Fig. 5e for 10 minutes, respectively, and then we compared the difference in fluorescence intensity between the compound in the plaques and in the background areas. We found that the transformation process of SHA-2 in plaques slower than SHA-2 in background, regardless of the exposure under green light. The difference is especially noticeable when using green light, which is also consistent with the wavelength range of excitation of DASAs (linear form). This difference means that many more SHA-2 outside the plaque are converted into **B** than those on the plaque. Taken together, our data suggested that Aβ species could change the isomerization process of SHA-2 in a biological relevant environment.

**Fig. 5.**
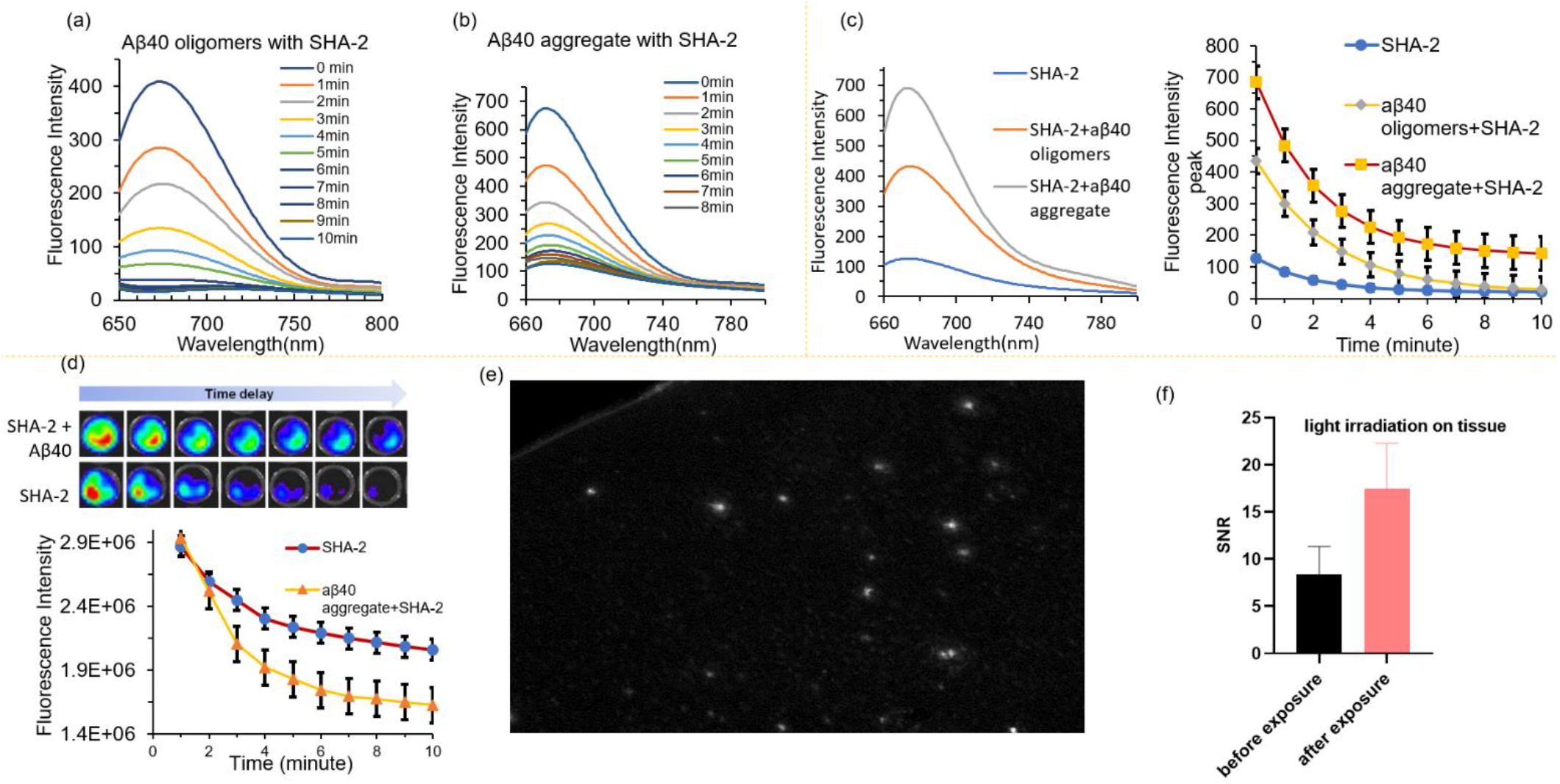
Aβ40 species to tailor DASAs switch. (**a**) Fluorescence of SHA-2 between 10 min in Aβ40 oligomers PBS solution; (**b**) Fluorescence of SHA-2 between 10 min in Aβ40 aggregate PBS solution; (**c**) Fluorescence comparison between SHA-2/SHA-2+ Aβ40 oligomers/SHA-2+Aβ40 aggregate at 0 min and Kinetics fitting results for different SHA-2 solution; (**d**) IVIS plate imaging of SHA-2 with or without Aβ40 species; (**e**) Microscopic images for SHA-2 stained with APP/PS1 mice brain slice; (**f**) Signal-to-Noise ratio (SNR) after green light exposure tissue for 10 minutes.

### 4. SHA-2 binding induced Aβ40 conformational change

Our results above suggested that Aβ species could change the isomerization process of SHA-2. On the other hand, it will be interesting to investigate whether the binding of SHA-2 to Aβ species can alter the process of structural change of Aβ peptides in solutions. To this end, we examined the secondary and tertiary structures of Aβ40-only and Aβ40-SHA-2 in water via molecular dynamic simulations. For Aβ40 in water, statistics were collected from five independent 400-ns simulations, while for SHA-2-Aβ40, the four 400-ns simulations with favorable binding were used. From the secondary structure analysis, Aβ40 monomer, known as an intrinsically disordered protein, mostly adopts a random coil structure (46 ± 5 %) with some bend (26 ± 5 %) and turn (15 ± 5 %) contents, consistent with experimental observations and previous simulations of Aβ40 monomer in aqueous solution,^41–42^ and the structural variation is relatively large from one simulation to another, indicated as large error bars in Fig. 6(a). The binding of SHA-2 has little effect on the secondary structure of Aβ40 monomer given the uncertainty of computed secondary structure content, while the tertiary structure of Aβ40 monomer is substantially altered by SHA-2. In Fig. 6(b), two representative Aβ40 configurations from MD snapshots of Aβ40 and Aβ40-SHA-2 systems are shown, and it is evident that the binding of SHA-2 leads to a more extended tertiary structure of Aβ40 monomer.

**Figure 6.**
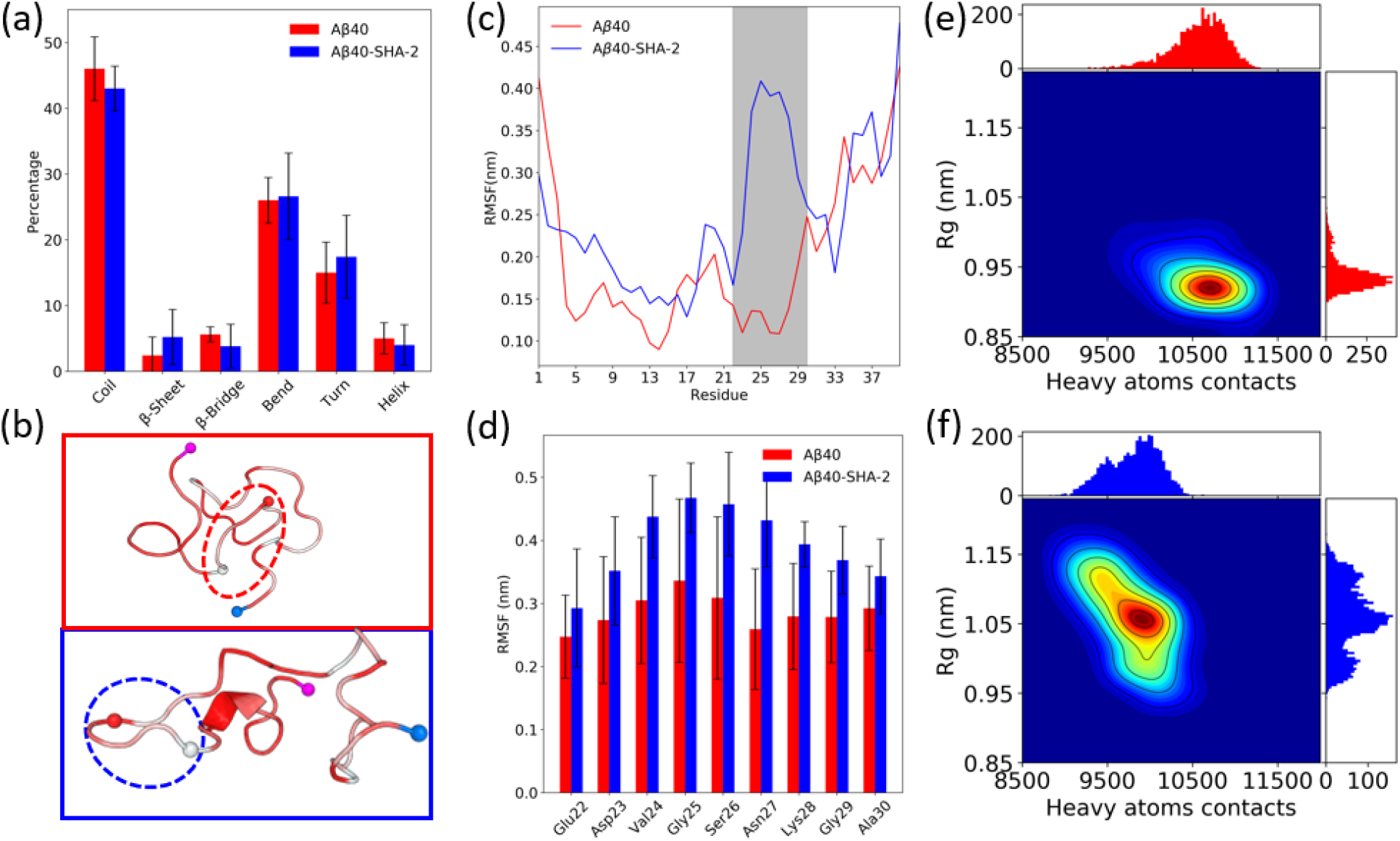
(**a**). Aβ40 secondary structure comparisons with the averages and error bars from all four simulations; (**b**). Representatives MD snapshots of Aβ40 monomer (blue box) and Aβ40-SHA-2 (red box). Protein color representation from white to red indicates the least hydrophobic to the most hydrophobic region, and the loop from Glu22 to Ala30 was circled with blue and red dashed lines for Aβ40 and Aβ40-SHA-2 systems, respectively. N and C termini are colored in marine and magenta; (**c**). Residue-specific RMSF for Aβ40 and Aβ40-SHA-2 complex. The gray-color shaded region represented the most fluctuated region on Aβ40 (Glu22 to Ala30) when bound with SHA-2; (**d**). The averaged RMSF among all four simulations where SHA-2 bound to Aβ40 at similar region for Glu22 to Ala30. The error bars represent the standard deviation of four simulations; (**e**) and (**f**): 2D histograms of heavy atoms contacts and radius of gyration for Aβ40 and Aβ40-SHA-2 systems, respectively.

To further analyze the tertiary structures of Aβ40 with and without bound SHA-2, residue-specific root mean square fluctuation (RMSF) were computed for Aβ40 and Aβ40-SHA-2, and the results from one 400-ns simulation were shown in Fig. 6(c). We find that the residues on the loop Glu22-Ala30 of Aβ40 exhibits significantly enhanced fluctuations upon the binding of SHA-2. As mentioned earlier, SHA-2 forms hydrogen bonds with Glu22 (or Asp23) and Lys28, which result in the loop Glu22-Ala30 more exposed to water as shown in **Fig. 6b**, due to the steric constraints imposed by hydrogen bonds. As a result of the unfavorable interactions between water and the hydrophobic residues (Val24, Gly25, Gly29, and Ala30), the loop region becomes floppier, giving rise to larger RMSFs. This tertiary structural change was observed in all four simulations where SHA-2 binds to the similar position on Aβ40 with favorable hydrophilic and hydrophobic interactions, and **Figure 6d** shows the averaged RMSF over all four simulations with standard deviations labelled as error bars. From MD simulation snapshots (**Figure 6c**), residues Glu22 to Ala30 in apo Aβ40 form a bend structure and collapse to other parts of peptide so that the region becomes less exposed to solvent, and the amphipathic feature of Aβ40 enables this region to stay folded and stable. However, the binding of SHA-2 to Aβ40 through the hydrogen bonds between SHA-2 and Glu22/Lys28 (or Asp23/Lys28) prevents the Glu22-Ala30 region from folding into peptide, and keeps it as a turn structure exposed to water with high RMSF during the course of 400 ns simulation. This simulation result is consistent with our SPEED dot-blotting studies, in which the binding of Aβs to its antibody 4G8 was weakened (**Fig. 3b**). 4G8 is a widely used antibody for immunological studies of Aβs, and its binding epitope on Aβ peptide is Leu17 to Val24, which falls in the binding segment of SHA-2 to Aβ peptides.

The enhanced exposure of the Glu22-Ala30 loop to water induced by the SHA-2 binding is expected to result in a reduction in the number of heavy atom contacts and an increase in the protein size of Aβ40. Panels (e) and (f) in **Fig. 6** display the 2D histograms of Aβ40 tertiary structures in terms of the number of heavy atom contacts and radius of gyration (Rg) for apo Aβ40 and Aβ40-SHA-2. Upon the binding of Aβ40, the number of heavy atom contacts decreases from 10572 ± 325 to 9803 ± 301, which can be attributed to the contact loss of the Glu22-Ala30 loop to other residues of Aβ40. Meanwhile, Rg increases from 0.94 ± 0.03 nm to 1.06 ± 0.05 nm. Both changes suggest that the binding of SHA-2 greatly affects the tertiary structure of Aβ40 monomer, but not its secondary structure. This observation was also found in the binding of some Aβ-aggregation inhibitors to Aβ40 monomer^43^, suggesting that SHA-2 might exhibit Aβ40 anti-aggregation activity.

### 5. Anti-aggregation experiments

The above results showed that SHA-2 (linear form) can bind with Aβ40 species and this prompted us to hypothesize that SHA-2 could inhibit Aβ40 monomer aggregation. In this regard, we incubated Aβ40 monomer without and with SHA-2 under dark, respectively. After 1 day, we run the SDS-PAGE Gel then silver staining. We found that in dark the SHA-2 can inhibit the Aβ40 monomer aggregation (**Fig.7a**). We also detect the amount of Aβ40 aggregate for these incubated solutions with Thioflavin T, which is the gold standard for detection of Aβ40 aggregate. The solution of Aβ40 incubated with SHA-2 under dark is significantly lower than the control incubated solution of Aβ40 without SHA-2 (*left,* **Fig. 7b**). To further verify our hypothesis, concentration-dependency experiments were carried out. The thioT fluorescence intensity of Aβ40 solution decreased with the increasing concentration of inhibitor SHA-2 (*Middle* and *right*, **Fig. 7b**). In addition, the peptides solutions were stained with ProteoStar, an amyloid-specific fluorescent dye. This reagent was previously shown to facilitate specific and sensitive detection of amyloid aggregates in living cells.^44^ Confocal microscopy were then employed to detect the presence of aggregates, the fluorescence intensity of the sample with SH-2 is significantly reduced, while the sample without SHA-2 can see obvious aggregates(**Fig. 7c**). These data indicated that SHA-2 could significantly inhibit the formation of aggregates.

**Figure 7.**
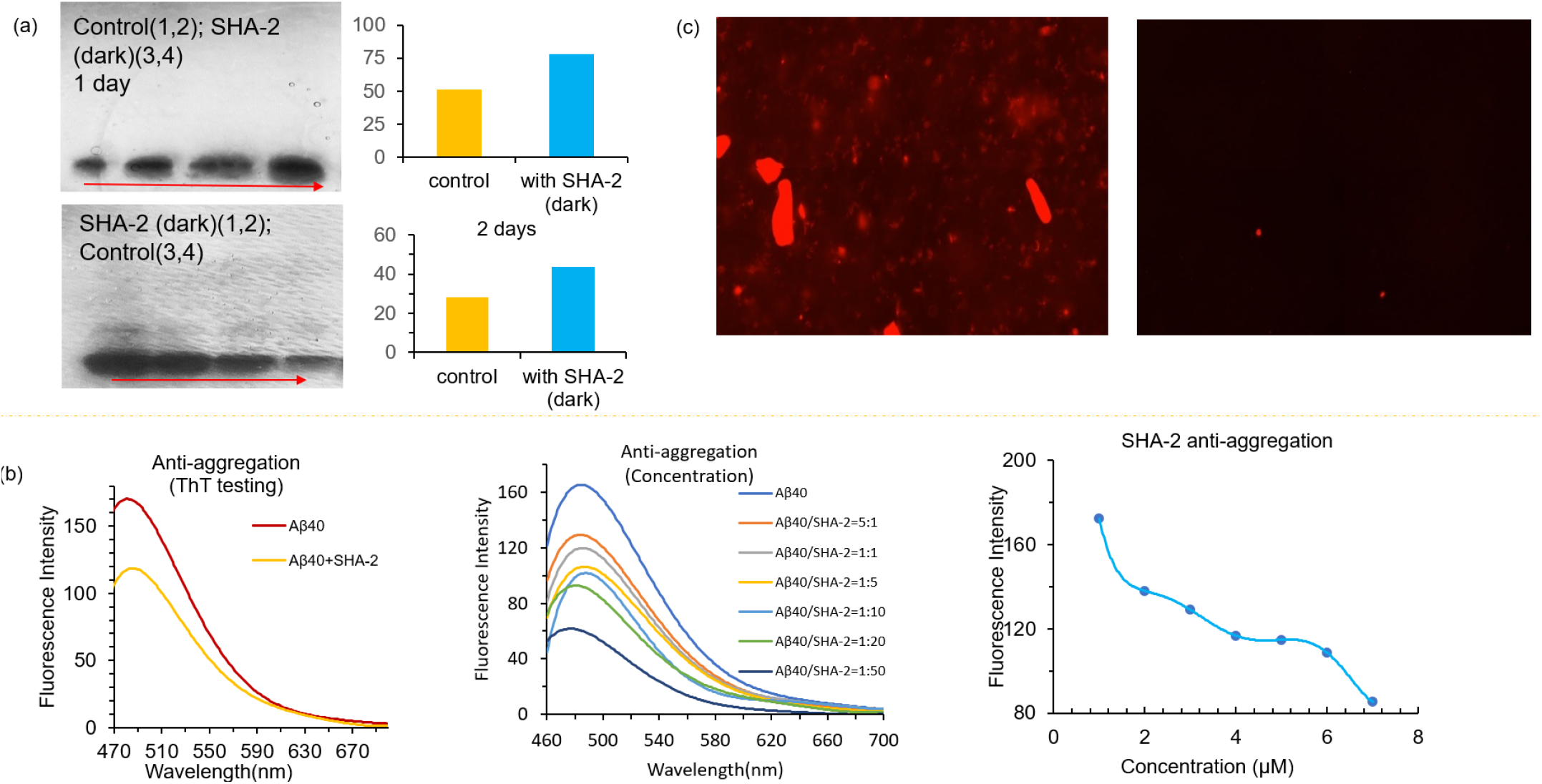
Anti-aggregation experiments. (a) Western Blot validation SHA-2 inhibited Aβ40 aggregation under dark condition (up: two samples on the left are Aβ40 without SHA-2, two samples on the right are Aβ40 with SHA-2; down: two samples on the left are Aβ40 with SHA-2, two samples on the right are Aβ40 without SHA-2; On the right is the comparison chart of the average gray scale of silver staining; (b) ThT fluorescence validation SHA-2 inhibited Aβ40 aggregation under different concentration; (c) Validation of SHA-2 inhibited Aβ40 aggregation with Proteostat ® aggresome detection kit.

## Conclusions and Discussions

In this report, we demonstrated that Aβ40 peptides could tailor the switching behavior of DASA compound SHA-2. In the presence of Aβ40 peptides, the conversion rate of linear-to-cyclic can be lowered. Reportedly, light or water can trigger the linear-to-cyclic isomerization of DASAs; however, the linear (open) form can be stabilized only in confined space of a water-soluble Pd^II^_8_ molecular vessel. Here, we reported the first peptides method to alter the switching property of DASAs in physiological environment. Our study signified that, in addition to light, peptides are also able to alter molecular switching of DASA compounds. Moreover, this tailoring of molecular switching process can be used to inhibit protein aggregation. In addition, our MD simulation results provided new ideas for developing inhibitors of Aβ aggregation.

The binding of SHA-2 to monomeric Aβ40 enhanced the structural favorability of SHA-2 to **A2** configuration. In the aqueous SHA-2 simulations, SHA-2 configurations were observed in all isomerization states including **A1, A2** and **A3**. The binding of SHA-2 to Aβ40 significantly enhanced the SHA-2 configuration towards **A2** configuration with hydrogen bond formations to peptide region Glu22-Lys28 which resulted in indoline moiety staying closer to more hydrophobic region around Phe19 that leads to a more stable **A2** configuration of SHA-2. The binding position also verified in our SPEED dot-blotting studies. The binding effect results in the increase of fluorescence intensity, prolong decay half-life of SHA-2, and slowing down of red-shift in Aβ40 monomer solution. All of our results indicated that the binding of SHA-2 with Aβ40 could slow down the linear-to-cyclic conversion rate.

The flexibility of peptide loop between Glu22 to Ala30 was significantly enhanced upon SHA-2 binding. For Aβ40 in solutions, we observed the region was in a stable bend structure but folded into peptide, so that the hydrophilic and hydrophobic parts from the region interacted favorably within peptide taking advantage of amphipathic character of Aβ40. Lazo *et al.* suggested that the region of Val24 to Lys28 could nucleate the intramolecular folding of the Aβ40 according to proton solution-state NMR.^45^ This indicates that our finding on monomeric Aβ40 in aqueous solution is consistent with experimental observations that the loop between Glu22 to Ala30 (Val24 to Lys28 was included in this region) was shown intramolecular folding with higher stability. When SHA-2 bound to Aβ40, the structure between Glu22 to Ala30 was maintained as turn configuration, however, the structure was observed to have high fluctuations as it stayed more exposed to solvent. Moreover, binding of SHA-2 to Aβ40 significantly reduced the heavy atoms contacts of protein, with an increased Rg, so that the protein exhibited less folded configuration. It implies that SHA-2 bound to protein resulted in different tertiary structure of protein and the peptide is less folded with higher fluctuation of region on Glu22 to Ala30, which suggested that SHA-2 is potentially able to inhibit the aggregation of Aβ40 by producing the off-pathway structures. This reductionof aggregation was verified with SDS-Page gel electrophoresis, fluorescence spectral tests, and Proteostat® aggresome detection kit.

The investigation of novel photoresponsivity molecules DASAs are still in their infancy. Our work shows that Aβ40 peptide can tailor molecular switch of the DASAs. Therefore, the role of Aβ40 in tailoring molecular switch of DASAs is encouraged to be understood from different perspective and needs further research. In addition to their responsive to visible or even NIR light, we envision that DASAs can be widely applied in biological system based on the peptides-tailor process, such as photo-therapeutics are most effective when activated by NIR and good tool to study protein folding/unfolding. Understanding the mechanism of the Aβ40 peptide tailor DASAs molecular switching behavior process is thus helpful for synthesizing a new DASA or apply to new peptide system in physiologically relevant environment.

