## supplemental for "β-Amyloid peptides tailor switching behaviors of Donor-Acceptor Stenhouse Adducts"

Tabel S 1 Computed internal energies, Gibbs free energies, and vertical emission wavelength of the three representative isomers of the linear SHA-2. The energies are reported relative to those of A1. ........ 4
Tabel S 2 Rate constants in min-1 and half time in min from fitting experimental fluorescence peak intensity of SHA-2 systems in PBS solution, as shown in Fig. SXX. ........ 5

### 1. Materials and Instruments

All chemicals were purchased through commercial vendors and used without further purification. Column chromatography was performed on a glass column slurry-packed with silica gel (60 Å, 40–63 mm; SiliCycle Inc.). Recombinant Aβ peptide (1-40) were purchased from rPeptide (A-1163-1). Aβ aggregates for in vitro studies were generated by slow stirring of Aβ40 in PBS buffer for 3 days at room temperature. $^{13}$C NMR and $^{1}$H NMR spectra were collected on a JOEL 500-MHz Spectrometer in $CDCl_3$, $CD_3OD$ or DMSO-d6 solutions at room temperature with tetramethylsilane (TMS, $\delta = 0$) as an internal standard. Absorption spectra were recorded on SpectraMax (Molecular Device, San Jose, CA). A fluorescence spectrophotometer F-7100 (Hitachi) was employed fluorescent spectra recording. LC-MS was carried out on an Agilent 1200 Series apparatus with an LC/MSD trap and Daly conversion dynode detector with UV detection at 254 nm. Tissue imaging was performed on Nikon Eclipse 50i. Two-photon imaging in vivo was conducted on an Olympus BX-51 microscope. Transgenic female 5xFAD mice and age-matched wild-type female mice were purchased from Jackson Laboratory. All animal experiments were approved by the Institutional Animal Use and Care Committee at Massachusetts General Hospital. An IVIS Spectrum animal imaging system (Perkin Elmer) was used for in vitro plate imaging. The IVIS Spectrum animal imaging system (PerkinElmer) was used for in vitro and vivo imaging. PROTEOSTAT® Protein aggregation imaging was taken by Zeiss Axio Imager Z2, Germany.

#### 1.1 Preparation of Aβ40 aggregates.

1.0 mg of Aβ40 peptide (TFA) was suspended in 1% ammonia hydroxyl solution (1.0 mL). Then 100 µL of the resulting solution was diluted 10-fold with PBS buffer (pH 7.4) and stirred at room temperature for 3 days. Transmission electron microscopy and Thioflavin T solution test were used to confirm the formation of aggregates [1].

#### 1.2 Preparation of Aβ40 oligomers.

The preparation was performed according to Kayed's reported procedure [2].

1.0mg of Aβ40 peptide (HFIP) was suspended in Hexafluoro-2-propanol (HFIP) solution (1.0 mL). Then 10 µL of the resulting solution was diluted 10-fold with PBS buffer (pH 7.4) and evaporated the HFIP at room temperature for 24 hours.

### 2. Synthesis and Fluorescence experiments

#### 2.1 Synthesis of SHA-2.

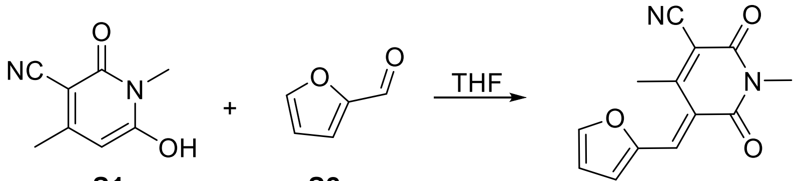

**(E)-5-(furan-2-ylmethylene)-1,4-dimethyl-2,6-dioxo-1,2,5,6-tetrahydropyridine-3-carbonitrile (S3).**

6-hydroxy-1,4-dimethyl-2-oxo-1,2-dihydropyridine-3-carbonitrile (S1) (500 mg, 3.04 mmol) and furfural S2 (307 mg, 3.04 mmol) was dissolved in 5 ml tetrahydrofuran, the mixture was stirred at room temperature. The reaction is monitored visually over the next 30 minutes. After the yellow color increase and the product begins to crash out. The reaction was allowed to stand for an additional 10 minutes. To this solution. Added 10 ml of a water solution (25% methanol) to the vial, shaken, filtered and rinsed twice with the same methanol/water mixture to yield a yellow solid (E)-5-(furan-2-ylmethylene)-1,4-dimethyl-2,6-dioxo-1,2,5,6-tetrahydropyridine-3-carbonitrile (S3) (375 mg, yield 51%). $^{1}$H-NMR (500 MHz, $CDCl_3$) δ 8.67 (s, 1H), 7.85 (s, 1H), 7.68 (s, 1H), 6.77 (s, 1H), 3.36 (s, 3H), 2.61(s, 3H). $^{13}$C-NMR (125 MHz, CDCl3) δ 162.0, 160.7, 158.4, 151.4, 150.7, 136.9, 128.7, 118.5, 115.9, 114.6, 105.2, 26.9, 19.1. HRMS: [M+Na]+ 265.0583, found 265.0589.

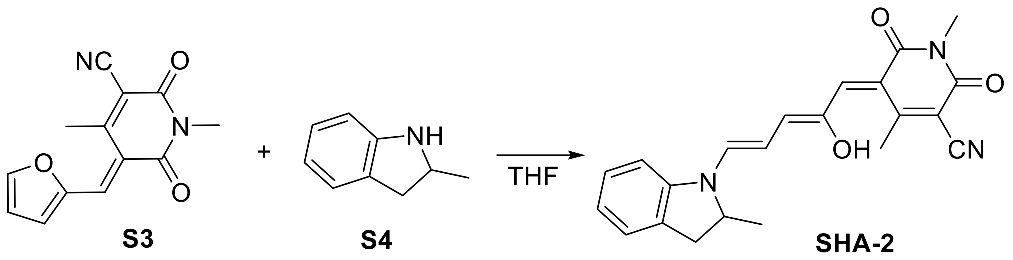

**(E)-5-((2Z,4E)-2-hydroxy-5-(2-methylindolin-1-yl)penta-2,4-dien-1-ylidene)-1,4-dimethyl-2,6-dioxo-1,2,5,6-tetrahydropyridine-3-carbonitrile (SHA-2).** The furan adduct S3 (100 mg, 0.41 mmol) was dissolved in 2 ml tetrahydrofuran and 2-methylindoline S4 (54 mg, 0.41 mmol) was added. The reaction mixture was stirred until starting material was consumed as observed by TLC. The reaction was then filtere and rinsed with diethyl ether to afford the product (70 mg, 46% yield). $^{1}$H-NMR (500 MHz, $CDCl_3$) δ 13.13 (s, 1H), 7.76 (d, J = 10 Hz, 1H), 7.33 (m, 2H), 7.23-7.21 (m, 1H), 7.14 (m, 1H), 6.84 (d, J = 10 Hz, 1H), 6.52-6.45 (m, 2H), 4.77 (m, 1H), 3.58-3.53 (m, 1H), 3.36 (s, 3H), 2.88 (d, J = 15 Hz, 1H), 2.43 (s, 3H), 1.47 (m, 3H). $^{13}$C-NMR (125 MHz, $CDCl_3$) δ 165.8, 162.3, 156.7, 150.9, 149.1, 144.9, 140.5, 135.9, 132.3, 128.8, 127.2, 126.9, 117.5, 111.1, 110.1, 107.4, 94.0, 58.3, 37.2, 36.4, 29.8, 27.5, 19.4. HRMS: [M+H]+ 376.1655, found 376.1651.

##### 2.2 Spectra of SHA-2 in $CH_2Cl_2$ or PBS solvents

The absorption spectrum of SHA-2 (10 μM) in the solution of PBS (10 mM, pH 7.4, 1% DMSO) or $CH_2Cl_2$ was recorded on SpectraMax Microplate Reader (Molecular Device, San Joe, CA). The SHA-2 stock solution was prepared in DMSO and all testing solutions contained a final concentration of 1% DMSO. The fluorescent spectra of SHA-2 (10 μM) in PBS (10 mM, pH 7.4, 1% DMF) or $CH_2Cl_2$ was performed on the F-7100 fluorescence spectrophotometer with Ex = 230 nm (in PBS), Ex = 630 nm (in DCM), slit 10/10, and 700 V for PMT.

##### 2.3 Fluorescence responses of SHA-2 in the presence of Aβ40 species

Fluorescence spectra of Aβ40 monomer (2.5 μM), Aβ40 oligomers (2.5 μM), Aβ40 aggregate (2.5 μM) in PBS (10 mM, pH 7.4, 1% DMSO) were recorded before and after the addition of SHA-2 (2.5 μM) with Ex = 630 nm, slit 10/10, and 700 V for PMT.

##### 2.4 In Vitro kinetic fluorescence spectral testing of SHA-2 with Aβ40 aggregates.

To test the interactions of **SHA-2** with Aβs, we used the following three-step procedure. In step **1**, 1.0 mL of double-distilled water or PBS (10 mM, pH 7.4) was added to a quartz cuvette as a blank control, and its fluorescence was recorded with the same parameters used for **SHA-2**. In step **2**, the fluorescence of an **Aβ40 species** solution (1.0 mL, 2.5 μM) was recorded with excitation at 630 nm and emission from 660 – 900 nm. In step 3, to the above **SHA-2** solution, 0.5 μL of Aβs (5 μM stock solution in DMSO) were added. Fluorescence readings from this solution were recorded as described in step **2**. And fluorescent spectra were recorded at different time points within 10 min (Ex = 630 nm, slit 10/10, 700 V for PMT). A blank control from step **1** was used to correct the final spectra from steps **2** and **3**.

##### 2.5 Epitope alteration experiment

Aβ monomers (1 μM, final concentration) was mixed with SHA-2 solution (2 μM, final concentration). After 1 h incubation, the mixture was spotted onto nitrocellulose membrane (0.2 μm, Bio-Rad) using 96-well Bio-Dot Microfiltration Apparatus. The membrane was blocked with 5% nonfat milk for 1 h and incubated with primary antibody (1:2000 diluted with 5% nonfat milk) at 4°C overnight with shaking. After washed with 0.1% Tween 20 in TBS (TBS-T) for 3 × 10 min, the membrane was incubated with HRP-conjugated secondary antibody (1:2000 diluted with 5% nonfat milk) for 1.5 h, followed by washing with TBS-T for 3 × 10 min. The membrane was then treated with enhanced chemiluminescent reagent (Pierce ECL Western Blotting Substrate) using an IVIS®Spectrum imaging system (Perkin Elmer, Hopkinton, MA) with blocked excitation filter and opened emission filter.

#### 3. Synthesis and Fluorescence experiments Study of SHA-2 inhibits aggregation of amyloid beta 40

##### 3.1 Fluorescence study SHA-2 anti-Aβ40 aggregation

**Solution prepared:**

Sample A: 2.5 μM of Aβ40 monomer solution in PBS (pH 7.4) incubated in room temperature under dark for 24 hours;

Sample B: 2.5 µM of Aβ40 monomer and 2.5 µM SHA-2 solution in PBS (pH 7.4) incubated in room temperature under dark for 24 hours;

Sample C: 2.5 µM of Aβ40 monomer and 2.5 µM SHA-2 solution in PBS (pH 7.4) incubated in room temperature under 400 W lamp for 24 hours;

Gradient study samples: A series of solutions containing 2.5 µM µM of Aβ40 monomer and various concentrations of SHA-2 (0.0, 0.5, 2.5, 12.5, 25, 50, 125 µM) incubated in room temperature under dark for 24 hours

Triplicate samples were prepared for each condition.

**Thioflavin T study SHA-2 anti-Aβ40 aggregation:**

10 µL of the above solution was added to 1.0 mL PBS solution. The resulting solution was then subjected to fluorescence spectrum recording with excitation = 430 nm and emission = 460-700 nm (baseline recording). To this solution, 10 µL of Thioflavin T (2.5 µM in PBS (pH 7.4)) was added, and the spectrum was recorded. The quantification was conducted at λem = 500 nm for Thioflavin T reading by subtracting the baseline reading.

##### 3.2 Western Blotting for SHA-2 with Aβ40 monomer.

To validate the anti-aggregation behavior of SHA-2, SHA-2 was incubated with Aβ monomers under light or dark condition for 24 h. The samples were subjected to gel electrophoresis (SDS-PAGE gel) and then stained with Pierce™ silver staining kit. The grayscale of each group was quantified by Image J software. Figure showed SHA-2 could only inhibit the aggregation of Aβ monomers under dark condition but not light condition. This may arise from the change of SHA-2 to the closed-form under light condition that cannot inhibit Aβ monomers.

##### 3.3 PROTEOSTAT® Protein aggregation imaging study

For sample A and sample B, according to PROTEOSTAT® Protein aggregation assay to quantitative detection of protein aggregates.

##### 3.4 Plate imaging of Aβ40 species with SHA-2.

The solution of Aβ40 aggregate (2.5 µM, 40 µL, in PBS, 10 mM, pH 7.4) and PBS were added to wells on a 96-well plate, specifically. The SHA-2 (5 µM, 20 µL) was added in and the final volume was 60 µL. Follow on the Plate images were recorded on the IVIS System. And images were recorded at different time points within 5 min.

##### 3.5 In vitro histological study.

A fresh brain tissue from a 24-month old APP/PS1 mouse was fixed in 4% formaldehyde for 24 hours and transferred into 30% sucrose at 4 °C until the tissue sunk. Then the tissue was embedded in OCT with gradual cooling over dry ice. The OCT embedded tissue block was sectioned into 25-µm slice with a cryostat. 25 M of SHA-2 in 20% ethanol/PBS was prepared as the staining solution. The brain slices were incubated with freshly prepared staining solution for 15 min at room temperature and then washed with 70% ethanol for 1min, 20% ethanol for 1min, followed by washing with double distilled water twice. Then the slice was covered with FluoroShield mounting medium (Abcam) and sealed with nail polish. Florescence images were obtained using the Nikon Eclipse 50i microscope with blue, green and red-light excitation channel, and then acquired florescence images again under green or red-light for 10 min, respectively.

#### 4. Quantum Chemical Calculations

We performed density functional theory (DFT) calculations to estimate the energetics of different isomers of linear SHA-2 in water, i.e., A1, A2, and A3 in Figure 1 of the main text. The geometries of isomers were minimized using M06-2X/6-31+G(d) with conductor-like Polarizable Continuum Model (CPCM) solvation model[3,4] for water, and vibrational analysis was performed to verify that the local minima were reached. The thermodynamic energies (i.e., internal energy, and Gibbs free energy) were computed at the same theory level within the harmonic approximation. M06-2X/6-31+G(d) has shown good performance in predicting the properties of other DASAs in previous studies[5,6,7,8]. To estimate the emission wavelengths, the first excited-state geometries of the isomers were optimized using time-dependent density functional theory (TDDFT) with M06-2X/6-31+G(d), and the vertical emission energy from first excited state to ground state was computed with the state-specific solvation correction.

The obtained results are summarized in Table S1. Consistent with the calculated results on other DASA molecules in previous studies, A1 isomer is the most stable one thermodynamically, and is anticipated to be the dominate isomer at ambient condition. Our TDDFT calculations suggest that the observed red shift may be attributed to the photo-induced isomerization from A1 to A2 or A3. Note that the calculated results are only meant to be qualitative, and an explicit quantum mechanical treatment of water may be needed for a more quantitative estimate.

Tabel S1 Computed internal energies, Gibbs free energies, and vertical emission wavelength of the three representative isomers of the linear SHA-2. The energies are reported relative to those of A1.

| Isomer | Internal energy (kcal/mol) | Gibbs free energy (kcal/mol) | Emission wavelength (nm) |
|---|---|---|---|
| A1 | 0 | 0 | 523 |
| A2 | 7.9 | 7.6 | 584 |
| A3 | 7.8 | 8.2 | 617 |

### 5. Kinetic Modeling of Fluorescence Data

The decay of the fluorescence intensity at about 680 nm is due to the gradual conversion of linear SHA-2 (A) to cyclic one (B), which has no emission around 680 nm. Therefore, we propose the following simplified kinetic model

$$A \underset{k_{-1}}{\overset{k_1}{\rightleftarrows}} B$$

First-order kinetics are assumed for both forward and backward reactions, and the time-dependent concentration of A is given by

$$\frac{[A](t)}{[A](0)} = \frac{k_1}{k}e^{-kt} + \frac{k_{-1}}{k},$$

Eq 1

where $k = k_1 + k_{-1}$, and $[A](t)$ is the concentration of A at time t. If we assume that the time dependence of fluorescence intensity follows that of concentration, we can fit the fluorescence peak intensity (around 680 nm) based on Eq. 1, and the fitting results for different solutions are shown in Fig. S4 and Table S2. Half time, define as $t_{1/2} = \ln 2 / k$, is also reported in Table S1. It is clear that upon the addition of Aβ40 to the PBS solution of SHA-2, the conversion of A to B is significantly hindered presumably due to the binding of SHA-2 to Aβ40.

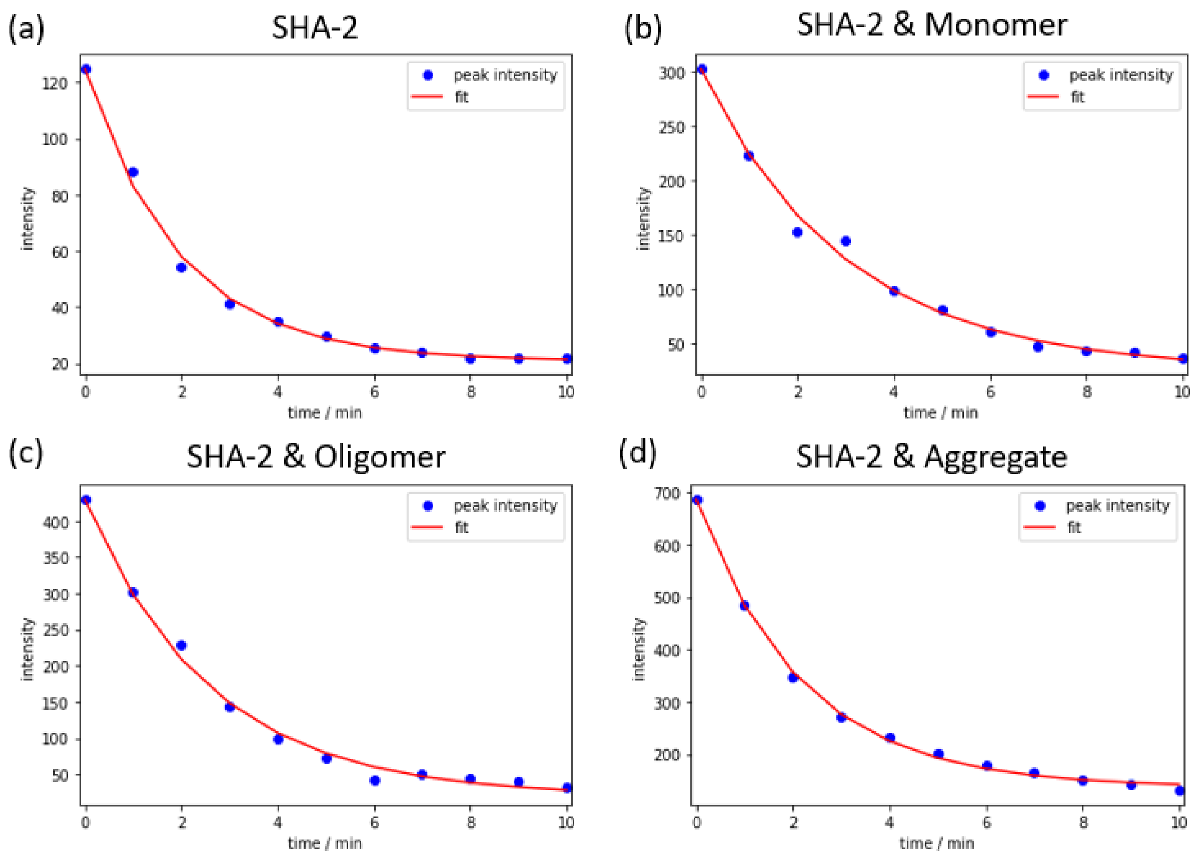

Fig S 1 Time-dependent fluorescence peak intensity from experiment (blue circles) and its fit from Eq. 1 (red lines) for (a) SHA-2 only, (b) SHA-2 and Aβ40 monomer, (c) SHA-2 and Aβ40 oligomers, and (d) SHA-2 and Aβ40 aggregate in PBS solution.

Tabel S 2 Rate constants in min-1 and half time in min from fitting experimental fluorescence peak intensity of SHA-2 systems in PBS solution, as shown in Fig. S1.

| System | $k_1$ ($\text{min}^{-1}$) | $k_{-1}$ ($\text{min}^{-1}$) | Half time (min) |
|---|---|---|---|
| SHA-2 | 0.428 | 0.085 | 1.35 |
| SHA-2 & Aβ40 monomer | 0.305 | 0.028 | 2.08 |
| SHA-2 & Aβ40 oligomer | 0.368 | 0.018 | 1.80 |
| SHA-2 & Aβ40 aggregate | 0.366 | 0.091 | 1.52 |

#### 6. Molecular Dynamics (MD) Simulations

MD simulations of SHA-2, Aβ40 monomer, and Aβ40 monomer bound to SHA-2 (denoted as Aβ40-SHA-2) in water were preformed independently. For each system, five independent simulations with different initial configurations and velocities were conducted using GROningen MAchine for Chemical Simulations (GROMACS) version 2018.4 package to improve statistical effciency.[9] The force field parameter of SHA-2 was generated using Automated forced field Topology Builder (ATB) version 3.0 web server.[10] Due to the novelty of the SHA-2, existing force field is not accessible, and ATB provided a good estimation for drug-like novel molecule based on GROMOS family force field (GROMOS 54a7)[11] in a GROMACS compatible format.[12,13] For consistency, Aβ40 monomer was also modeled with GROMOS 54a7 force field. Several benchmark studies suggested that GROMOS 54a7 gives a good alpha-beta-coil balance consistent with the experimental solution NMR study on Aβ40 monomer, and the calculated NMR chemical shifts, J-couplings, and radius of gyration for Aβ40 monomer and oligomers agree reasonably with experimental estimates.[14,15] It should be noted that MD force fields, in general, have inherent limitations and biases, and a recent study by Man and co-workers suggests that GROMOS 54a7 may be biased toward β-sheet and lead to incorrect kinetics when used for studying amyloid formation,[16] while other force fields may excessively stabilize helical structure.[17] SPC water model was used as the default water model for GROMOS 54a7.[18]

#### 6.1 MD simulation for SHA-2 in water

Single SHA-2 molecule was solvated in a cubic water box filled with roughly 2,171 SPC water models for all five simulations. Additional ions, 6 Na+ and 6 Cl-, were added to match experimental ion concentration 0.15 M. The solvated system first went through 5000 steps of energy minimization using steepest descent (SD) method followed by two equilibration processes. The first equilibration was performed under NVT ensemble for 500 ps using Berendsen thermostat with a coupling constant of 0.1 ps to maintain the temperature at 300 K.[19] The second equilibration was preformed using NPT ensemble for 20 ns using the v-rescale thermostat with a coupling constant of 0.5 ps and Berendsen barostat with a coupling constant of 1.0 ps to maintain the temperature and pressure at 300 K and 1 bar respectively.[19,20] The production simulation was then conducted under NPT ensemble with all hydrogen atoms being constrained using LINCS for 400 ns under the same thermostat and barostat as those in the second equilibration run.[21] All simulations were applied three-dimensional periodic boundary conditions with integration time step of 2 fs. The neighbor list was cut off at 1.2 nm using Verlet scheme.[22] Long-range electrostatic interactions were calculated with the smooth particle mesh Ewald method with a cutoff of 1.2 nm, non-bonded van der Walls interactions was using the plain cut off at distance of 1.2 nm.[23,24]

#### 6.2 MD simulation for Aβ40 monomer in water

The monomer Aβ40 was constructed from a completely extended conformation of the full-length peptide with free amine and carboxyl group placed at N and C termini, respectively. The side-chain protonation states was evaluated at pH 7.4 (experimental conditions) using H++ web server.[26–28] Aβ40 was first solvated in a cubic box with 147,036 SPC water containing 0.15 M of NaCl, and additional 3 Na+ ions were added to balance the charge. First, the solvated system went through the same minimization procedure discussed above, and the energy minimized system underwent a 1-ns MD simulation at high temperature (700K) under the NVT ensemble with the Berendsen thermostat (0.1ps coupling constant) to collapse each extended system. A similar protocol was employed by Rosenman *et al.* for obtaining collapsed configuration from extended structure.[29] All five independent collapsed configurations were generated and applied for Aβ40 and Aβ40-SHA-2 in aqueous simulations (see next subsection).

From the resulting collapsed configuration, a new simulation box was constructed with roughly 13,336 SPC water molecules and 0.15M NaCl concentration with 3 additional Na+ ions for neutralizing the charges. All systems underwent the same simulation protocol discussed above with slightly different equilibration procedure. Equilibrations were also conducted in two phases. For the first part, NVT ensemble was performed for 500 ps at 300 K with the Berendsen thermostat (0.1ps coupling constant), but additional position restraints were added on all heavy protein atoms. For second equilibration, NPT ensemble was applied with all restraints removed, v-rescale thermostat and Berendsen barostat with coupling constants of 0.5ps and 1ps to maintain the system temperature and pressure at 300 K and 1bar, respectively, and protein and solvent (including ions) were coupled separately. Production run was conducted for 400 ns using the same NPT ensemble. A representative snapshot of Aβ40 monomer is shown in Fig. S2. We note that as Aβ40 monomer is known to be an intrinsically disordered protein, there are relatively large secondary structural variation among the five independent simulations, as shown in Fig. 3b in the main text.

#### 6.3 MD simulation for Aβ40 monomer bound with SHA-2 complex (Aβ40-SHA-2)

To construct Aβ40-SHA-2 system, a similar procedure to that for Aβ40 monomer in water was adopted. The five collapsed Aβ40 configurations were employed and solvated in water, and then one SHA-2 molecule was randomly inserted into each system.[30, 31] All other MD simulations parameters were the same as those for Aβ40 monomer in water.

SHA-2 bound stably with Aβ40 monomer at similar positions in four out of five independent 400-ns Aβ40-SHA-2 simulations (in the other simulation, SHA-2 weakly bound to the N-terminus of Aβ40, exposing itself to water), and a representative MD snapshot of SHA-2 binding to Aβ40 is shown in Fig. S5(b). SHA-2 formed two hydrogen bonds (HBs) with Glu22 (or Asp23) and Lys 28 of Aβ40. The hydroxyl group near dioxo-tetrahydropyridine of SHA-2 served as a HB donor to Glu22-Oε. Note that one of these four simulations with stable binding showed that SHA-2 can also possibly serve as hydrogen bond donor to Asp23-Oδ, but HB between SHA-2 and Glu22-Oε or Asp23-Oδ were mutually exclusive, namely that once HB is formed between SHA-2 and Glu22, it is unlikely to form HB with Asp23, and vice versa. The carbonyl oxygen on dioxo-tetrahydopyridine ring served as HB acceptor in the HB with positively charged Lys28-Nζ. In addition to HBs, the binding of SHA-2 to Aβ40 is further stabilized by the pi-pi interaction between the indoline of SHA-2 and Phe19 of Aβ40.

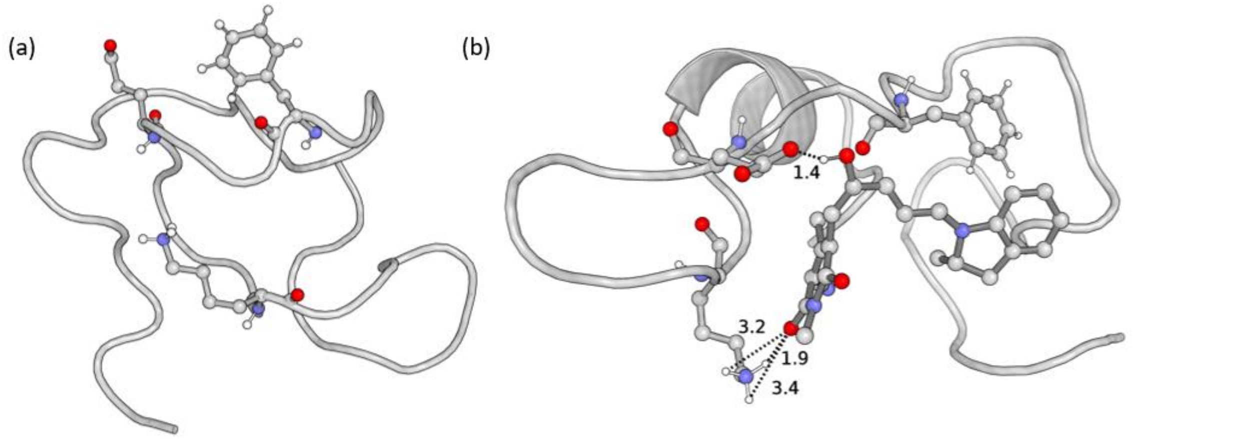

Fig S 2 Representative MD snapshots for (a) Aβ40 monomer, and (b) Aβ40 monomer bound with SHA-2 (Aβ40-SHA-2) in PBS solution. SHA-2 binding position on Aβ40, Glu22 and Lys28 are labelled and possible hydrogen bonds formed between Aβ40 and hydroxypyridone of SHA

#### 7. Binding of SHA-2 to Aβ40 fibril predicted by molecular docking

SHA-2 binding position on Aβ40 fibril (PDB code: 6SHS) was predicted using affinity-based method provided by AutoDock (version 4.2.6)1. SHA-2 conformation was allowed to be flexible with 4 rotatable bonds during docking to obtain the optimal binding positions. AutoGrid4 was used for establishing grid maps for interaction energies that was further used by AutoDock for docking.[32] The grid map was set to cover the entire Aβ40 fibril with grid dimension to be 68 Å × 120 Å × 40 Å and grid spacing parameter to be 0.625 Å. For docking calculations, genetic algorithm (GA) was applied to search for docking positions with 50 independent GA runs and a population size of 1500, and the maximum number of evaluations was capped at 2,500,000. The most stable binding position is shown in Fig. S6.

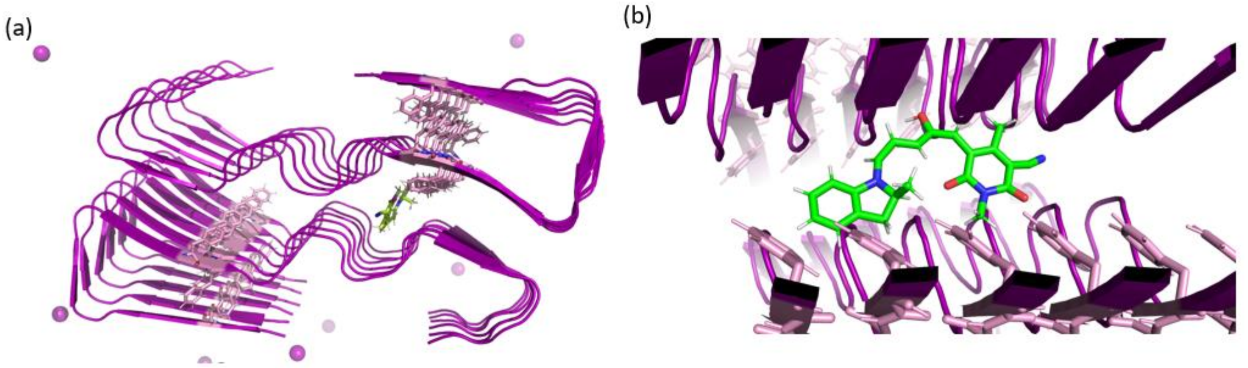

Fig S 3 (a) Binding position of SHA-2 on Aβ fibril (PDB code: 6SHS) predicted from docking; (b) the zoom-in view of binding site.

(6) Donato, M. D.; Lerch, M. M.; Lapini, A.; Laurent, A. D.; Lagatti, A.; Bussotti, L.; Lhrig, S. P.; Medved, M.; Jacquemin, D.; Szymański, W.; Buma, W. J.; Foggi, P.; Feringa, B. L.

Shedding Light on the Photoisomerization Pathway of Donor-Acceptor Stenhouse Adducts, *J. Am. Chem. Soc.* **2017**, *139*, 15596.
(7) Mallo, N.; Iranmanesh, H.; Kennedy, A. D. W.; Luis, E. T.; Ho, J.; Harper, J. B.; Beves, J. E. Structure-function relationships of donor-acceptor Stenhouse adduct photochromic switches, *Chem. Sci.*, **2018**, *9*, 8242.
(8) Lerch, M. M.; Donato, M. D.; Laurent, A. D.; Medved', M.; Lagatti, A.; Bussotti, L.; Lapini, A.; Buma, W. J.; Foggi, P.; Szymański, W.; Feringa, B. L. Solvent Effects on the Actinic Step of Donor-Acceptor Stenhouse Adduct Photoswitching, *Angew. Chem. Int. Ed.* **2018**, *57*, 8063.
(9) Abraham, M. J.; Murtola, T.; Schulz, R.; Páll, S.; Smith, J. C.; Hess, B.; Lindah, E. Gromacs: High Performance Molecular Simulations through Multi-Level Parallelism from Laptops to Supercomputers. *SoftwareX* **2015**, *1–2*, 19–25. https://doi.org/10.1016/j.softx.2015.06.001.
(10) Stroet, M.; Caron, B.; Visscher, K. M.; Geerke, D. P.; Malde, A. K.; Mark, A. E. Automated Topology Builder Version 3.0: Prediction of Solvation Free Enthalpies in Water and Hexane. *J. Chem. Theory Comput.* **2018**, *14*, 5834.
(11) Schmid, N.; Eichenberger, A. P.; Choutko, A.; Riniker, S.; Winger, M.; Mark, A. E.; van Gunsteren, W. F. Definition and Testing of the GROMOS Force-Field Versions 54A7 and 54B7. *Eur. Biophys. J.*, **2011**, *40*, 843.
(12) Malde, A. K.; Zuo, L.; Breeze, M.; Stroet, M.; Poger, D.; Nair, P. C.; Oostenbrink, C.; Mark, A. E. An Automated Force Field Topology Builder (ATB) and Repository: Version 1.0. *J. Chem. Theory Comput*., **2011**, *7*, 4026.
(13) Koziara, K. B.; Stroet, M.; Malde, A. K.; Mark, A. E. Testing and Validation of the Automated Topology Builder (ATB) Version 2.0: Prediction of Hydration Free Enthalpies. *J. Comput. Aided. Mol. Des*. **2014**, *28*, 221.
(14) Gerben, S. R.; Lemkul, J. A.; Brown, A. M.; Bevan, D. R. Comparing Atomistic Molecular Mechanics Force Fields for a Difficult Target: A Case Study on the Alzheimer's Amyloid β-Peptide. *J. Biomol. Struct. Dyn*. **2014**, *32*, 1817.
(15) Somavarapu, A. K.; Kepp, K. P. The Dependence of Amyloid-β Dynamics on Protein Force Fields and Water Models. *ChemPhysChem* **2015**, *16*, 3278.
(16) Man, V. H.; He, X.; Derreumaux, P.; Ji, B.; Xie, X.-Q.; Nguyen, P. H.; Wang, J. Effects of All-Atom Molecular Mechanics Force Fields on Amyloid Peptide Assembly: The Case of Aβ 16–22 Dimer. *J. Chem. Theory Comput*. **2019**, *15*, 1440.
(17) Best, R. B.; Buchete, N.-V.; Hummer, G. Are Current Molecular Dynamics Force Fields Too Helical? *Biophys. J.* **2008**, *95*, L07.
(18) Berendsen, H. J. C.; Postma, J. P. M.; van Gunsteren, W. F.; Hermans, J. Interaction Models for Water in Relation to Protein Hydration.; Pullman, B., Ed.; Reidel, Dordrecht, *Intermolecular Forces*, **1981**, *14*, 331.
(19) Berendsen, H. J. C.; Postma, J. P. M.; van Gunsteren, W. F.; DiNola, A.; Haak, J. R. Molecular Dynamics with Coupling to an External Bath. *J. Chem. Phys*. **1984**, *81*, 3684.
(20) Bussi, G.; Donadio, D.; Parrinello, M. Canonical Sampling through Velocity Rescaling. *J. Chem. Phys*. **2007**, *126*, 014101.
(21) Hess, B. P-LINCS: A Parallel Linear Constraint Solver for Molecular Simulation. *J. Chem. Theory Comput*. **2008**, *4*, 116.
(22) Páll, S.; Hess, B. A Flexible Algorithm for Calculating Pair Interactions on SIMD Architectures. *Comput. Phys. Commun*. **2013**, *184*, 2641.
(23) Essmann, U.; Perera, L.; Berkowitz, M. L.; Darden, T.; Lee, H.; Pedersen, L. G. A Smooth Particle Mesh Ewald Method. *J. Chem. Phys*. **1995**, *103*, 8577.
(24) Darden, T.; York, D.; Pedersen, L. Particle Mesh Ewald: An N ·log( N ) Method for Ewald Sums in Large Systems. *J. Chem. Phys*. **1993**, *98*, 10089.
(25) Kabsch, W.; Sander, C. Dictionary of Protein Secondary Structure: Pattern Recognition of Hydrogen-Bonded and Geometrical Features. *Biopolymers* **1983**, *22*, 2577.
(26) Gordon, J. C.; Myers, J. B.; Folta, T.; Shoja, V.; Heath, L. S.; Onufriev, A. H++: A Server

for Estimating PKas and Adding Missing Hydrogens to Macromolecules. *Nucleic Acids Res*. **2005**, *33 (Web Server)*, W368–W371.
(27) Myers, J.; Grothaus, G.; Narayanan, S.; Onufriev, A. A Simple Clustering Algorithm Can Be Accurate Enough for Use in Calculations of PKs in Macromolecules. *Proteins Struct. Funct. Bioinforma*. **2006**, *63*, 928.
(28) Anandakrishnan, R.; Aguilar, B.; Onufriev, A. V. H++ 3.0: Automating PK Prediction and the Preparation of Biomolecular Structures for Atomistic Molecular Modeling and Simulations. *Nucleic Acids Res*. **2012**, *40 (W1)*, W537–W541.
(29) Rosenman, D. J.; Connors, C. R.; Chen, W.; Wang, C.; García, A. E. Aβ Monomers Transiently Sample Oligomer and Fibril-Like Configurations: Ensemble Characterization Using a Combined MD/NMR Approach. *J. Mol. Biol.* **2013**, *425*, 3338.
(30) Zhang, S.; Iwata, K.; Lachenmann, M. J.; Peng, J. W.; Li, S.; Stimson, E. R.; Lu, Y. -a.; Felix, A. M.; Maggio, J. E.; Lee, J. P. The Alzheimer's Peptide Aβ Adopts a Collapsed Coil Structure in Water. *J. Struct. Biol.* **2000**, *130*, 130.
(31) Lazo, N. D.; Grant, M. A.; Condron, M. C.; Rigby, A. C.; Teplow, D. B. On the Nucleation of Amyloid β-Protein Monomer Folding. *Protein Sci*. **2009**, *14*, 1581.
(32) Morris, G. M.; Ruth, H.; Lindstrom, W.; Sanner, M. F.; Belew, R. K.; Goodsell, D. S.; Olson, A. J. Software News and Updates AutoDock4 and AutoDockTools4: Automated Docking with Selective Receptor Flexibility. *J. Comput. Chem*. **2009**, *30*, 2785.

### 9. NMR Spectra of key compounds

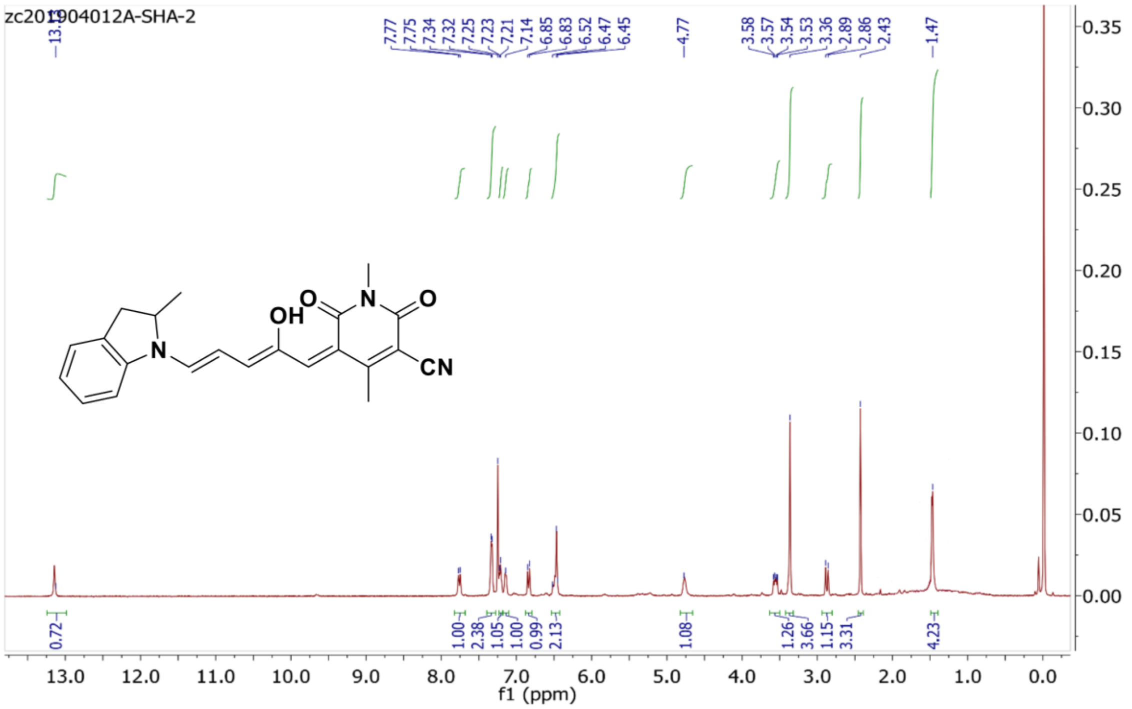

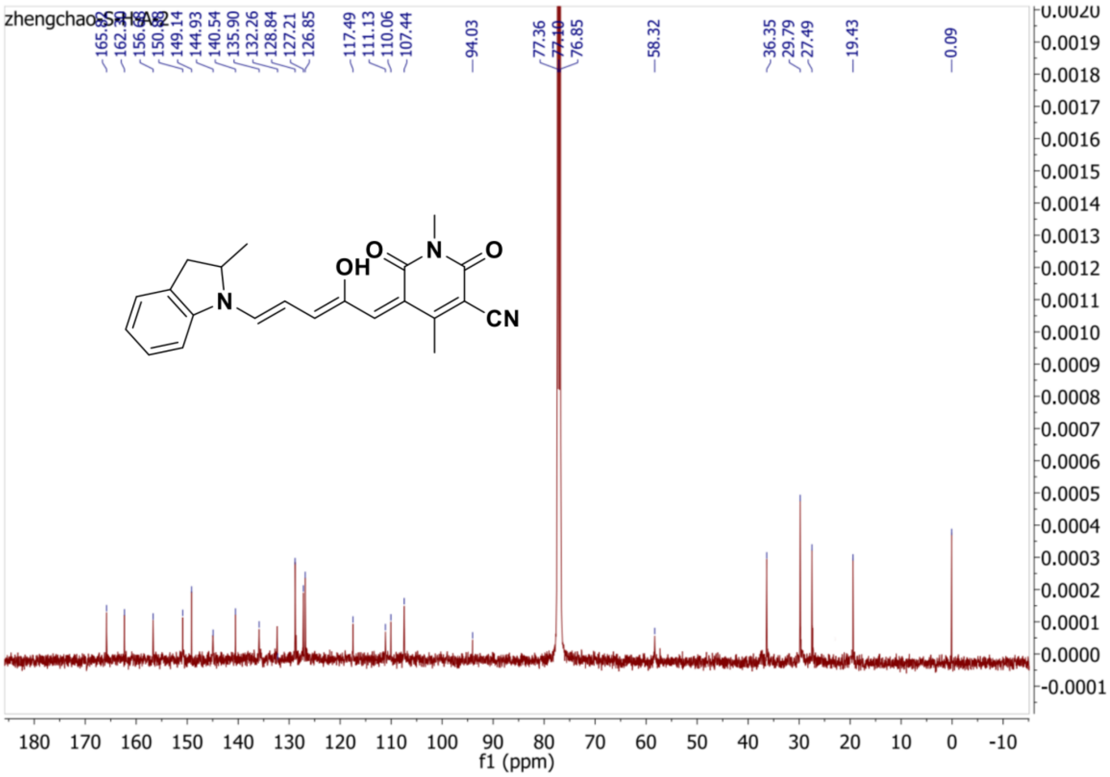

Fig S 5 NMR spectra of compound SHA-2
